# Making Proteomics Accessible: RokaiXplorer for interactive analysis of phospho-proteomic data

**DOI:** 10.1101/2023.08.22.553639

**Authors:** Serhan Yılmaz, Filipa Blasco Tavares Pereira Lopes, Daniela Schlatzer, Marzieh Ayati, Mark R. Chance, Mehmet Koyutürk

## Abstract

In this work, we present RokaiXplorer, an intuitive web tool designed to address the scarcity of user-friendly solutions for proteomics and phospho-proteomics data analysis and visualization. RokaiXplorer streamlines data processing, analysis, and visualization through an interactive online interface, making it accessible to researchers without specialized training in proteomics or data science. With its comprehensive suite of modules, RokaiXplorer facilitates the analysis of phosphosites, proteins, kinases, and gene ontology terms. The tool offers functionalities such as data normalization, statistical testing, enrichment analysis, subgroup analysis, report generator, and multiple visualizations, including volcano plots, bar plots, heatmaps, tables, and interactive networks. Additionally, RokaiXplorer allows researchers to effortlessly deploy their own data browsers, fostering the sharing of research data and findings interactively. By providing simplicity, efficiency, and multi-level data analysis within a single application, it is poised to become a valuable resource for the scientific community working with phospho-proteomic data. Access RokaiXplorer at: http://explorer.rokai.io

## 1 Introduction

In the field of proteomics and phospho-proteomics, there is a growing need for user-friendly tools that enable researchers to analyze and visualize data with minimal training. To address this need and make proteomics data analysis easily accessible to researchers without expertise in computer and data sciences, we introduce RokaiXplorer, a comprehensive framework for performing exploratory analysis on proteomic and phosphorylation data in an interactive environment (Figure 1).

**Figure 1.**
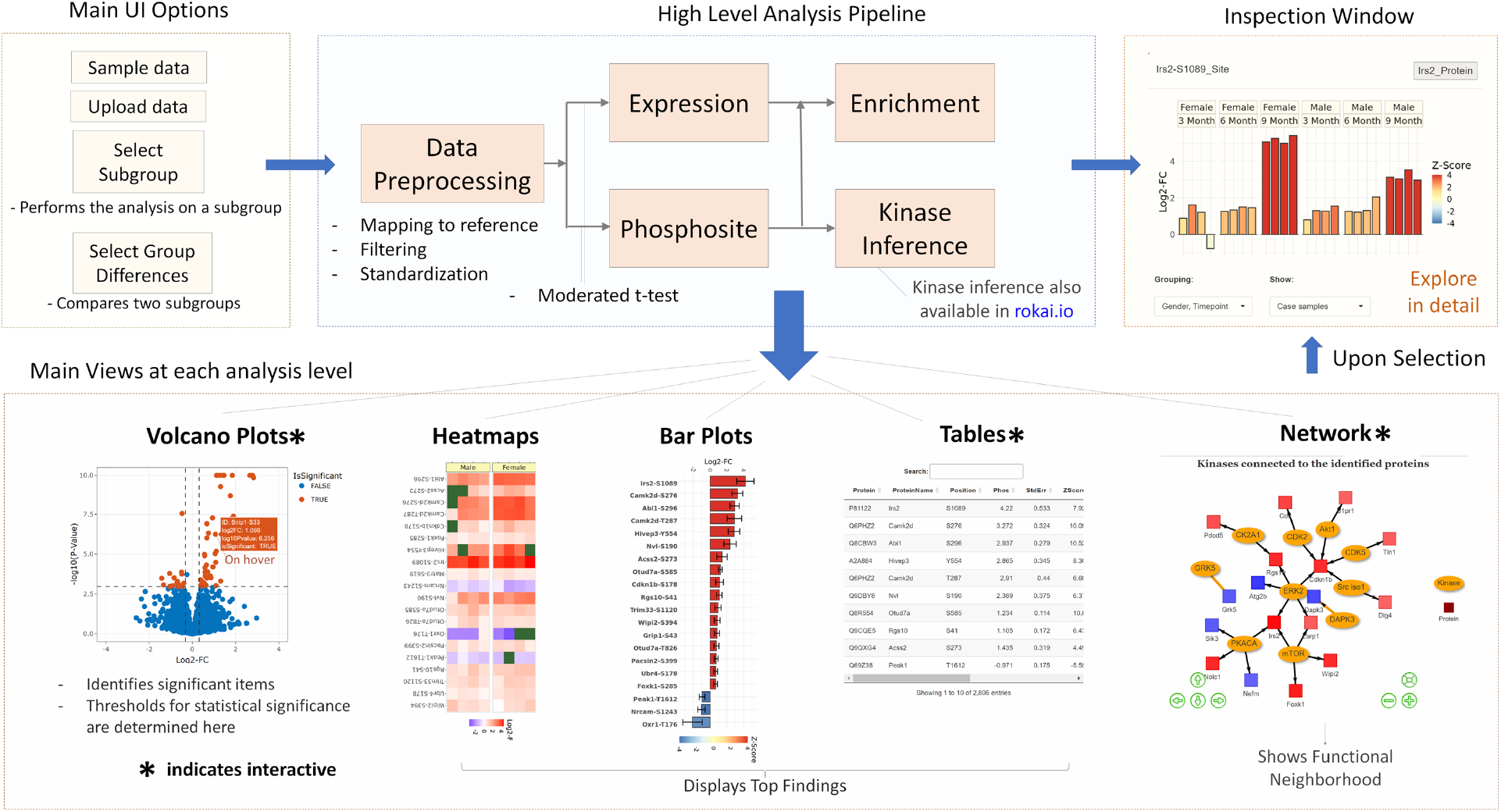
The workflow and key idea of RokaiXplorer.

RokaiXplorer offers a range of functionalities that operate at five levels: Phosphosite, Phospho-protein, Protein expression, Pathway Enrichment, and Kinases. It allows for the identification of significant dysregulation at each level and presents the top findings through various visualizations, including volcano plots, heatmaps, bar plots, tables, and a network view. One of the distinguishing features of RokaiXplorer is its interactivity, which enables users to click on selected items in the visualizations to access an Inspection Window. This window provides comprehensive information about the selected items, including the source of evidence for dysregulation, quantifications and raw data for all samples.

Getting started with RokaiXplorer is straightforward and user-friendly. The application provides an interactive tutorial that guides users through the initial steps, making it easy to familiarize themselves with the tool. To begin using RokaiXplorer, users only need two types of input data: quantification data and meta data. The quantification data is a .csv file that contains the phosphorylation levels of each phosphosite/peptide, with each row representing a specific site and the columns containing quantification values for multiple samples. The meta data file complements the quantification data by providing additional information about the samples, such as their grouping. The main group field, which is mandatory, specifies the case/control status of the samples, while optional additional groups can be utilized to focus the analysis on specific subgroups if desired. RokaiXplorer also supports the input of protein expression data, enhancing its versatility for comprehensive analyses.

### 1.1 Main features

RokaiXplorer offers a comprehensive suite of modules that enable researchers to perform various analyses on their datasets (Figure 2). One of its key functionalities is dataset normalization, which ensures accurate and reliable comparisons between samples. By normalizing the data, RokaiXplorer minimizes potential biases and enhances the statistical power of subsequent analyses.

**Figure 2.**
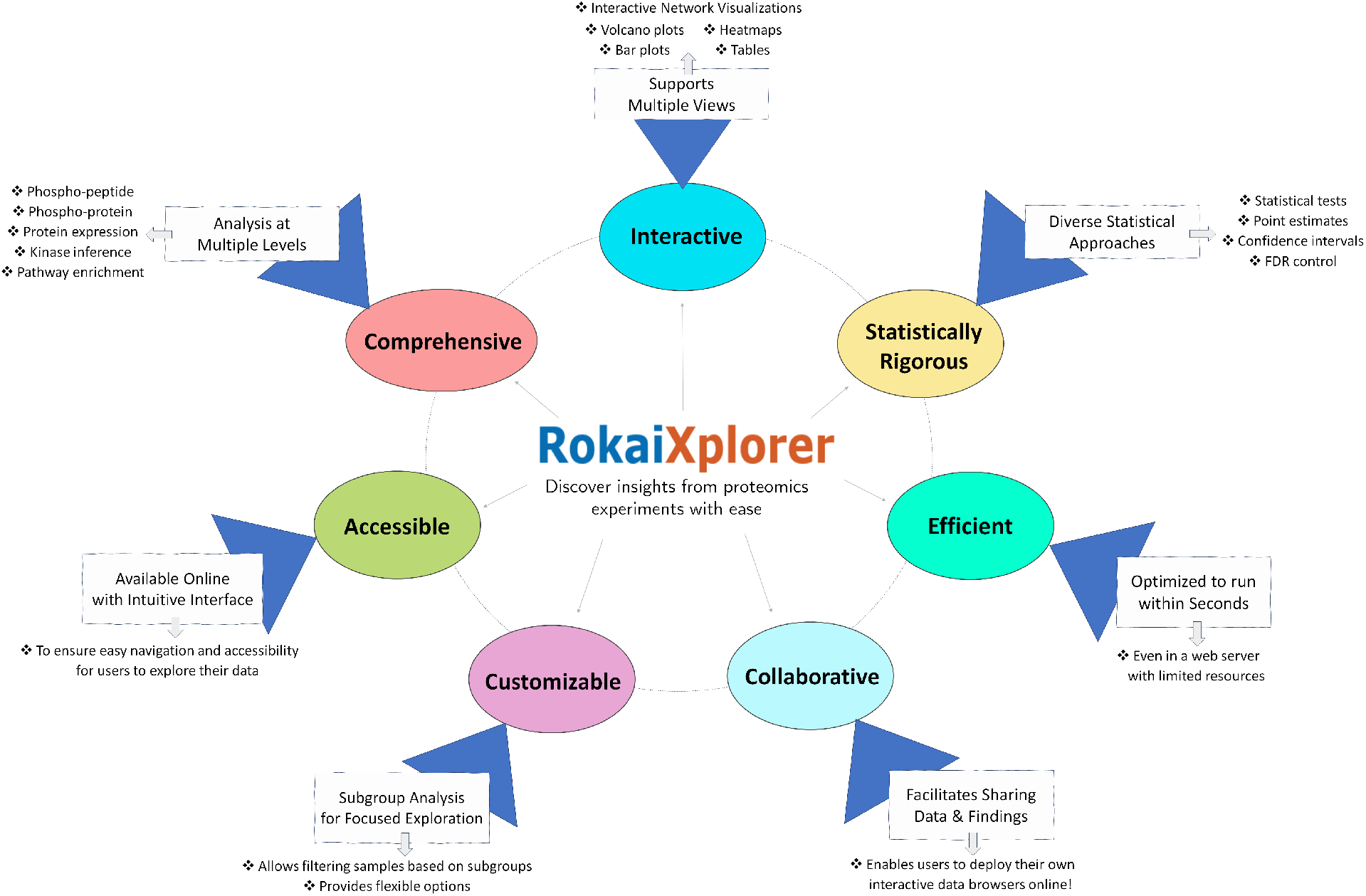
The main features of RokaiXplorer.

In addition to dataset normalization, RokaiXplorer facilitates the identification of dysregulated proteins, peptides, and potential biomarkers. It employs statistical tests, such as moderated t-tests, to assess dysregulation with precision, aiding in the identification of molecular signatures associated with specific conditions or diseases. Additionally, RokaiXplorer extends its analysis beyond individual proteins by providing insights into kinase activities. Through the utilization of the RoKAI algorithm, it infers kinase activities based on observed dysregulation patterns of phosphosites, contributing to a deeper understanding of cellular processes and signaling pathways.

To uncover the biological context of the dysregulated phospho-proteomic profiles, RokaiXplorer incorporates enrichment analysis of gene ontology (GO) terms. By identifying over-represented biological processes and molecular functions, researchers can gain valuable insights into the functional implications of their data. The GO enrichment analysis is performed using a chi-squared test with Yate’s correction, ensuring reliable and statistically significant results.

In addition, RokaiXplorer goes beyond traditional data analysis tools by offering a robust Report Generator feature. This feature simplifies the analysis of data for multiple subgroups and streamlines the process of exporting results as formatted Excel tables. With the Report Generator, researchers can effortlessly investigate the impact of variables such as gender or tissue type by performing separate analyses for each subgroup of interest. By selecting the desired analysis type and defining grouping variables, users can generate customized reports with a single click. This convenient and user-friendly functionality enhances the efficiency of data analysis and facilitates the dissemination of research findings.

### 1.2 Customization options in RokaiXplorer

Customization options in RokaiXplorer go beyond simple data processing and analysis. The tool provides researchers with the ability to tailor their analysis to specific subgroups and species, allowing for a more targeted investigation. By utilizing these options, researchers can focus their analysis on particular subpopulations or target organisms of interest (Figure 3).

**Figure 3.**
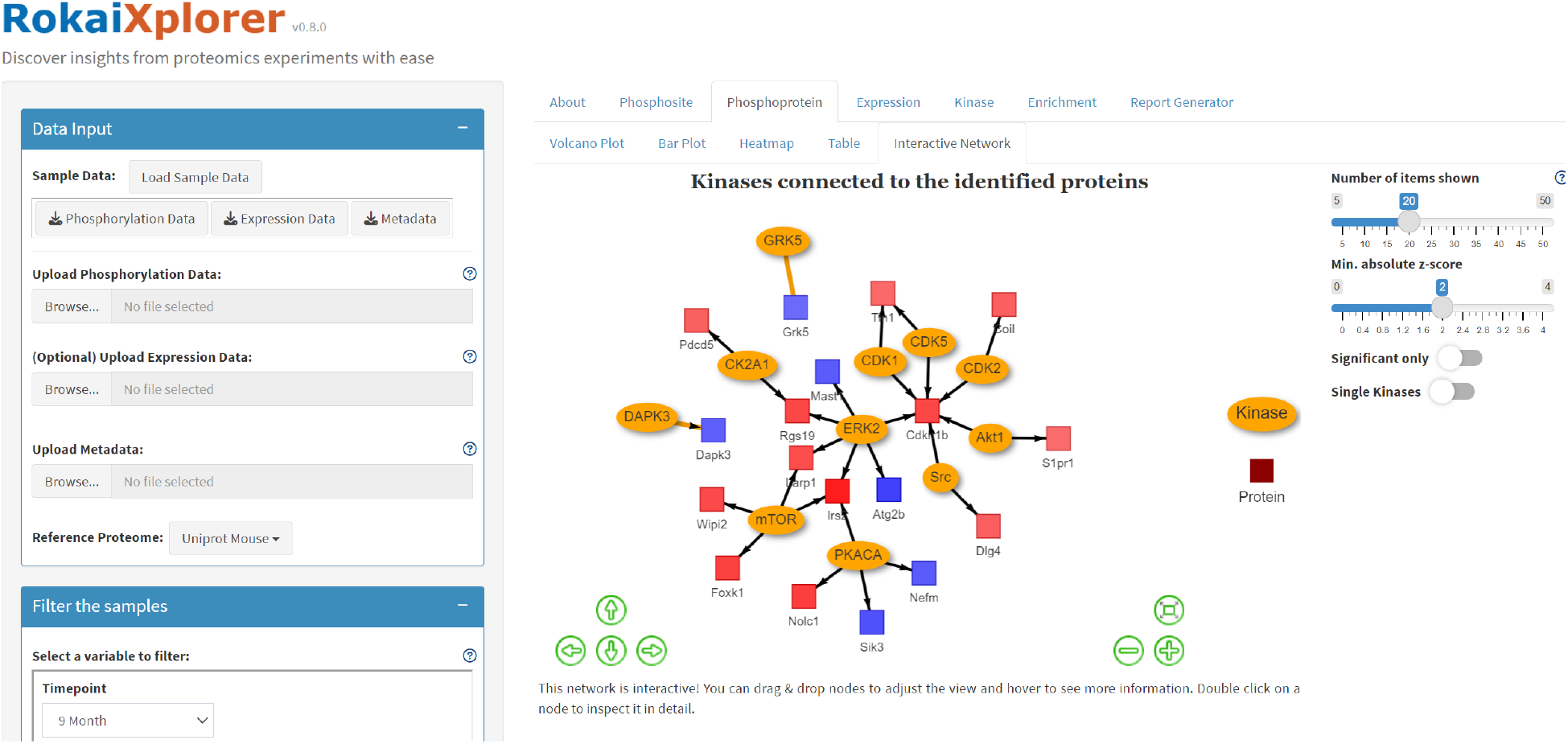
A snapshot of the user interface for RokaiXplorer as of v0.8.0. The options to select specific subgroup or species are shown on the left. The active tab in the figure displays the results of interactive network visualization for phospho-protein analysis.

One of the customization features is the ability to filter the analysis based on specific subgroups. For instance, users can select a subgroup such as “Gender *→* Male” to restrict the analysis to male samples only. By filtering the dataset in this manner, researchers can obtain results that are specific to the chosen subgroup, allowing for subgroup-specific insights and comparisons.

Additionally, RokaiXplorer supports multiple reference proteomes, accommodating various species of interest. The tool currently provides reference proteomes for Human (Homo sapiens), Mouse (Mus musculus), and Rat (Rattus norvegicus). Researchers working with proteomic data from these species can leverage the respective reference proteomes to enhance the accuracy and relevance of their analyses. This species-specific customization ensures that the results obtained from RokaiXplorer are aligned with the biological context and characteristics of the target organisms.

Furthermore, RokaiXplorer enables researchers to explore group differences between two sub-groups. This functionality is particularly valuable for comparative studies, where researchers want to investigate the patterns or dysregulation profiles between two specific groups. By selecting the desired subgroups for comparison, users can gain a deeper understanding of the molecular distinctions and uncover potential biomarkers or targets that are unique to each subgroup.

### 1.3 Interactive data browser: Share your discoveries feature

To foster collaboration and knowledge sharing, RokaiXplorer offers the “Share Your Discoveries” feature, which enables researchers to deploy their own interactive applications showcasing their data and analysis results. With this feature, the applications can be accessed online with the user data and settings already pre-loaded, allowing collaborators and other researchers to explore the data and gain valuable insights. This feature can enhance the impact of proteomic discoveries and facilitates interdisciplinary collaborations.

Deploying RokaiXplorer with preloaded input data is a straightforward process. Researchers can easily prepare and deploy their applications by following a few steps using the provided R scripts in the Github repository ^1^. These steps include installing R, RStudio, and Rtools (for Windows users), creating an RStudio project, downloading the RokaiXplorer source code, and installing the required R libraries. Once the setup is complete, researchers can run RokaiXplorer in deployment mode, customize the application for their specific data and configuration, and make modifications to the application’s title, descriptions, and about page. Additionally, RokaiXplorer allows users to export configuration files, enabling them to set desired analysis parameters for the online application and ensure reproducibility of results. Finally, researchers can deploy their application to shinyapps.io, a popular platform for hosting and sharing Shiny applications. By setting up a shinyapps.io account, connecting it to RStudio, and deploying the application, researchers can freely and effortlessly share their interactive RokaiXplorer application with others through a unique link, making their findings accessible to a wider audience.

### 1.4 Example application on Alzheimer’s disease

We utilized the capabilities of RokaiXplorer to analyze proteome and phospho-proteome data from a mouse hippocampus tissue study on Alzheimer’s disease (AD) [Yilmaz et al., 2023]. The dataset includes variables such as time (3, 6, and 9 months), sex (male and female), and genetic background (5XFAD versus wild type), which correspond to specific AD phenotypes such as Aβ42 plaque deposition, memory deficits, and neuronal loss. The goal of this study was to explore temporal and sex-linked variations in AD, focusing on biomarker discovery and identification of potential clinical targets.

This study involved various analyses to understand the phosphoproteome changes in the hippocampus of 5XFAD mice during Alzheimer’s Disease progression. We investigated the temporal and sex-linked patterns in phosphorylation, aiming to estimate the disease burden and identify trends over time, followed by a statistical analysis to identify specific dysregulated phosphopeptides between the case (5XFAD) and control (wildtype) mice groups. In addition, we compared phosphorylation patterns to protein expression levels to assess the complementarity of phosphorylation to protein expression. In addition, we identified consistent phosphoproteins that could potentially serve as markers for Alzheimer’s Disease, and investigated regulatory mechanisms involved in phosphorylation events through kinase inference analysis. Finally, an enrichment analysis was conducted to understand the biological pathways and networks impacted by the observed phosphoproteome changes. Together, these analyses provided a multi-faceted approach to uncovering the complex dynamics of phosphorylation and its implications in Alzheimer’s Disease progression. To facilitate the interpretation of our findings and promote free exploration of the data and results by other researchers, we utilized the RokaiXplorer application to develop the interactive tool AD-Xplorer. The findings are presented in the form of a live data browser with intact analysis capabilities, which can be accessed online at: https://yilmazs.shinyapps.io/ADXplorer

In the browser, the analysis can further be specified to focus on a particular subgroup. For example, selecting “9 Month” and “Female” on the left panel displays the findings for that group by performing the analysis after filtering the samples that fit the criteria. All presented capabilities are made generic and can be readily applied on other datasets abd studies, including the deployment of the dataset online as a live browser. The groups to customize the analysis are specified in a metadata file and deploying a dataset online only requires a configuration file (that can be generated via the online interface), a markdown file (to specify the descriptions on the front page) and the input data files. RokaiXplorer supports data from all proteomics quantification methods (e.g., label-free, SILAC, isobaric labeling).

Overall, we anticipate that RokaiXplorer will be an appreciated tool in the community to analyze phospho-proteomic data because of its simplicity and speed, enabling the analysis of data at different levels in one application. RokaiXplorer is available at: http://explorer.rokai.io

## 2 Methods

### 2.1 Data input

The input data required for RokaiXplorer consists of two data files and one additional data file which is optional. The following provides detailed information on the data formats for each file:

#### Phosphorylation Data

The phosphosite quantification data should be provided in CSV format. The file should contain the following columns:

- **Protein (first column):** This column should contain the Uniprot protein identifier.
- **Position (second column):** This column specifies the position of the modified phosphosite on the protein.
- **Samples (multiple columns):** Each column represents a sample, and the values in each column indicate the phosphorylation intensity of the corresponding phosphosite for that sample. The intensities should not be log-transformed, as this step is performed within the application.

#### Metadata

The metadata file should also be in CSV format and contain the following information:

- **RowName (first column):** This column provides the name of the group specifier.
- **Samples (multiple columns):** Each column represents a sample, and the values in each column indicate the group identity for that sample.
- **Group (first row):** This row is necessary and specifies the main group that determines the case/control status of the samples.
- **Other Groups (multiple rows):** You can use the optional rows in the metadata file to specify additional groups for the samples. These additional groups allow you to filter the samples and focus the analysis on a particular subgroup of interest.

Please ensure that your metadata file is in CSV format. The main group, which determines the case/control status, is required, while other group specifications are optional.

#### Expression Data (Optional)

If available, you can include protein expression data in CSV format. The file should have the following columns:

- **Protein (first column):** This column contains the Uniprot protein identifier.
- **Samples (multiple columns):** Each column represents a sample, and the values in each column indicate the expression intensity of the corresponding protein for that sample. The intensities should not be log-transformed, as this step is performed within the application.

Including protein expression data is optional, but if provided, it should be in CSV format.

### 2.2 Data preprocessing

#### 2.2.1 Notation

Let ***V*** *∈* ℝ^*n×m*^ denote the input data matrix for phosphorylation, where the rows denote phosphosites and the columns denote the samples, and let *V* [*i, j*] refer to an entry of this matrix corresponding to phosphosite *i* and sample *j*. Let 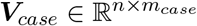 and 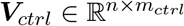 denote the subsets of this matrix correspond to case and control samples respectively, having *n* phosphosites, *m*_*case*_ case samples and *m*_*ctrl*_ control samples.

#### 2.2.2 Optional step: Filtering samples for a subgroup

If desired, the users have the option to filter the samples (columns) to narrow down the analysis to a specific subgroup. This step is carried out before any other analysis steps. By doing this, only the data related to the selected subgroup will be used for the analysis. It is essentially equivalent to excluding the data for other subgroups from the input altogether.

#### 2.2.3 Log-transformation and normality assumption

As a first step of preprocessing, we apply a log transformation on the quantification matrix ***V*** to make sure that it approximately follows a normal distribution:

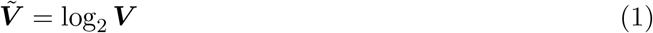

where 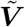 the matrix after the transformation. After the transformation, we assume that each column ***v***[:, *j*] of 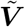 follows a normal distribution *𝒩* (*µ*_*j*_, *σ*_*j*_).

#### 2.2.4 Optional step: Centering for variance stabilization

As an optional step after the log transformation, we center each sample (column) to have 0 mean value by substracting the sample means 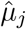:

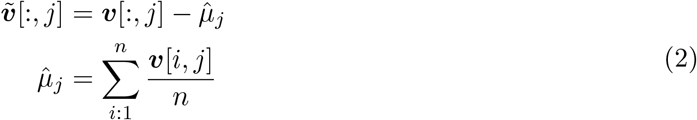

Note that, the missing values are omitted during the computation of the sample mean 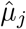. This step is enabled by default and is recommended to balance out potential systematic differences that may occur between the samples

### 2.3 Statistical inference at phosphosite level

As a first to identify phosphosites that are significantly different between the case and control samples, we compute the fold changes. Let *q*[*i*] denote the log fold change for phosphosite *i*, which is equal to the following:

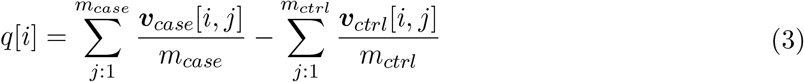

#### 2.3.1 Optional step: Centering the fold changes

As alternative approach to balance out any potential systematic bias between the case and control groups, the option to center the log fold changes are provided by subtracting the mean across all phosphosites:

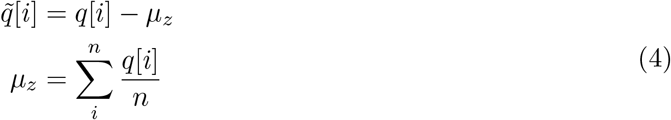

#### 2.3.2 Statistical tests

To determine the statistical significance of the log fold changes ***q***, we consider various models that differ in how the standard errors are estimated.

##### Z-test

The first and the simplest option is to estimate the standard errors based on the standard deviation across the phosphosites and perform a z-test based on this.

Let *s*_*z*_ be sample standard deviation of ***q*** across all phosphosites. Here, if we assume the standard error *σ*[*i*] of each phosphosite to be the same and equal to *s*_*z*_, the z-score *z*[*i*] for each phosphosite follows a normal distribution:

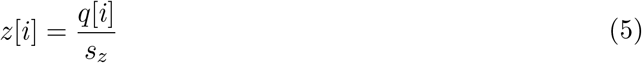

Based on the inverse normal distribution and the z-scores ***z***, the corresponding p-values are computed to the statistical significance. In addition, Benjamini-Hochberg [Benjamini and Hochberg, 1995] procedure is applied to limit the false discovery rate (FDR) of the findings.

Note that, this test is the simplest option with the weakest assumptions. It should only be applied in cases where the number of samples for each (case or control) group are too low and the standard deviations cannot be measured accurately (e.g., when is only a single sample). Otherwise, a t-test is more appropriate.

##### Pooled t-test

The second option is to estimate the standard error based on the standard deviation across the samples and perform a pooled t-test based on this.

Let *s*_*case*_[*i*] and *s*_*ctrl*_[*i*] be sample standard deviations estimated across the samples (columns) for each phosphosite *i*, and let *s*_*pooled*_[*i*] denote the pooled standard deviation for phosphosite *i* which is given as follows:

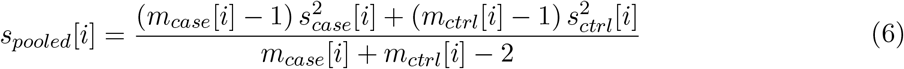

where *m*_*case*_[*i*] and *m*_*ctrl*_[*i*] represent the number of case/control samples with quantifications (i.e., having non-missing data) for phosphosite *i*. Assuming normality, independence between the samples, and equal variances between two groups (i.e., case and control), the t-statistic *t*[*i*] for phosphosite *i* follows a t-distribution with degrees of freedom *df* [*i*] such that:

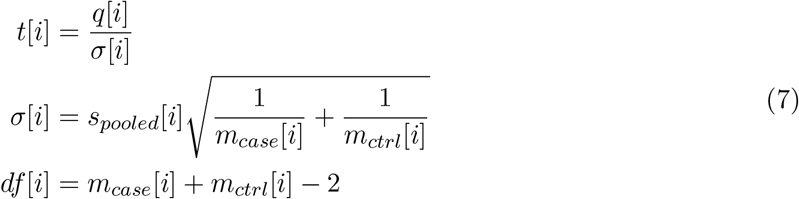

and the statistical significance and p-values are computed accordingly based on the t-distribution. Note that, this test requires at least *m*_*case*_ *≥* 2 and *m*_*ctrl*_ *≥* 2 to be performed.

##### Moderated t-test

If desired, as a potential improvement to pooled t-test, the moderated t-test [Smyth, 2004] can be performed, which utilizes an empirical Bayes method to shrink the pooled sample variances towards a common value and to augment the degrees of freedom for the individual variances. For this purpose, we utilize the implementation in limma package of R [Ritchie et al., 2015]. Specially, we utilize the *SquuezeVar* function which takes the pooled standard deviations *s*_*pooled*_[*i*] as input and returns the moderated standard deviations *s*_*mod*_[*i*] and the extra degrees of freedom gained *df*_*ext*_. Based on these, the moderated t-test is performed:

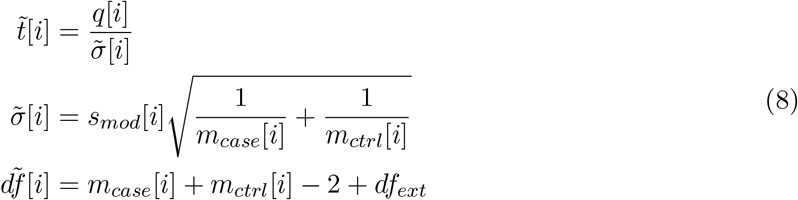

where the moderated t-statistic 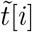 follows a t-distribution having 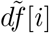 degrees of freedom.

##### Defaults

By default, if there are at least *m*_*case*_ *≥* 2 and *m*_*ctrl*_ *≥* 2 samples, a moderated t-test is performed. If that is not case, or if the analysis is to performed for a single sample (e.g., for heatmaps), a z-test is performed.

To ensure generalization and to simplify notation in the following sections, we will adopt the assumption that a t-test is performed. Additionally, we will consider the z-test as a special case of the t-test, where the parameters *σ*[*i*] are represented as *s*_*z*_ and the degrees of freedom *df* [*i*] are treated as *∞*.

### 2.4 Statistical inference at phospho-protein level

After assessing the significance of phosphosites, we combine their results to perform statistical inference at the protein level, again comparing the case samples with the control samples. Let *q*[*i*] be the resultant log2 fold change, and *σ*[*i*], *df* [*i*] be the corresponding standard error and degrees of freedom obtained from t-test for phosphosite *i*.

To perform the inference at the protein level, we first compute the mean log-fold changes *q*_*p*_[*j*] for each protein *j*:

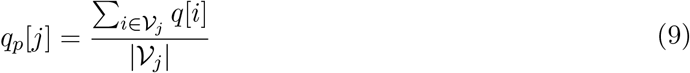

where *𝒱*_*j*_ denotes the set of phosphosites corresponding to protein *j*.

To estimate the pooled standard error *σ*_*p*_[*j*] and the corresponding degrees of freedom *df*_*p*_[*j*] in the estimation of the mean log-fold changes for each protein *j*, we use the Satterthwaite approximation [Satterthwaite, 1946]:

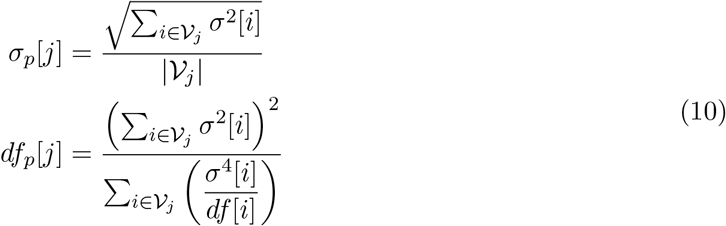

Based on these estimations, to compute the significance of a protein *j*, a t-test is performed with the t-statistic *t*_*p*_[*j*]:

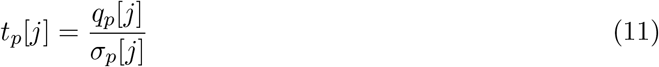

which follows a t-distribution with *df*_*p*_[*j*] degrees of freedom under the null hypothesis.

### 2.5 Optional: Statistical inference for protein expression

The statistical inference for protein expression follows the same methodology employed in the phosphosite level analysis, which includes pooled/moderated t-tests or z-tests as described in the *Statistical inference at phosphosite level* section. The key difference is that, in this case, the analysis is performed at the protein level instead of the phosphosite level.

### 2.6 Statistical inference at kinase level

We use the notation ***W***_*ks*_ to represent the kinase-substrate network, which consists of interactions between *n*_*kin*_ kinases and *n* phosphosites. We obtain this network from either the PhosphoSitePlus [Hornbeck et al., 2015] or Signor [Licata et al., 2020] databases. Typically, this network is sparse, with a value of 1 in the entry *w*_*ks*_[*i, j*] indicating that kinase *i* targets phosphosite *j*.

To identify dysregulated kinases that exhibit significant differences between case and control samples, we employ two approaches for inferring kinase activities. The first approach is a simple one, involving the calculation of mean substrate phosphorylation. This approach considers the phosphorylation (log-FC) of the known targets of a kinase and takes the mean value as the inferred activity of that kinase. In contrast, the RoKAI algorithm is a more comprehensive approach that utilizes a functional network to enhance the accuracy and robustness of kinase activity inference [Yilmaz et al., 2021].

#### 2.6.1 Mean substrate phosphorylation (without RoKAI)

To perform the kinase activity inference based on the mean substrate phosphorylation, we first compute the mean log-fold changes *q*_*k*_[*i*] for each kinase *i*:

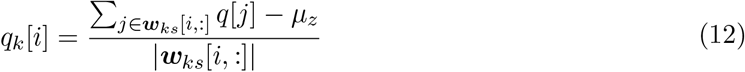

where ***w***_*ks*_[*i*, :] denotes the set of phosphosites that are known targets of kinase *i* and *µ*_*z*_ denotes the mean log fold change across all phosphosites (see Equation (4)).

To estimate the pooled standard error *σ*_*k*_[*i*] and the corresponding degrees of freedom *df*_*k*_[*i*] in the estimation of the mean log-fold changes for each kinase *i*, we use the Satterthwaite approximation [Satterthwaite, 1946]:

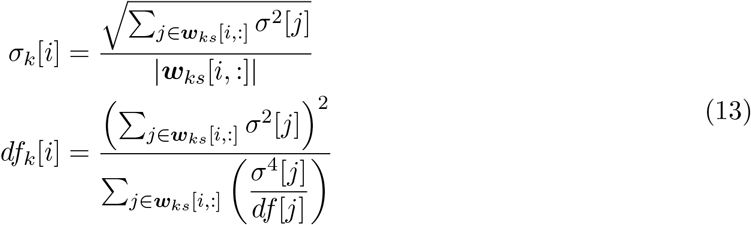

Based on these estimations, to compute the significance of a kinase *j*, a t-test is performed based on the t-statistic *t*_*k*_[*i*]:

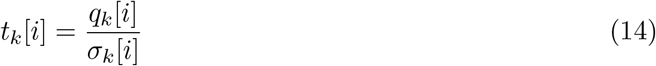

which follows a t-distribution with *df*_*k*_[*i*] degrees of freedom under the null hypothesis.

#### 2.6.2 Inference of kinase activities using RoKAI algorithm

The RoKAI algorithm [Yilmaz et al., 2021] is a method that propagates phosphorylation levels in a functional network. The network includes kinase-substrate associations, protein-protein interactions between kinases, and structure distance and co-evolution evidence for interactions between phosphosites. Using an electric circuit model, the algorithm transfers node potentials through the network using a conductance matrix ***C***. By solving a linear system, the algorithm computes node potentials, which enables the propagation of phosphorylation levels. The algorithm then infers kinase activities based on mean phosphorylation of known targets of the kinase using the propagated values.

Since RoKAI algorithm employs a linear model, the inferred activity of a kinase can be expressed as a weighted summation of phosphorylation levels (i.e., log fold changes) of known targets of a kinase, along with other phosphosites in the kinase’s functional neighborhood. In this section, we discuss the statistical methods used to determine the significance of the inferred kinase activities based on RoKAI. The following section cover the process of obtaining the weights that express the underlying formula in the RoKAI algorithm.

Let ***W*** denote the weighting matrix between *n*_*kin*_ kinases and *n* phosphosites, where *w*[*i, j*] represents the weight of phosphosite *j* in the inferred activity *q*_*a*_[*i*] of kinase *i* in the RoKAI inference such that:

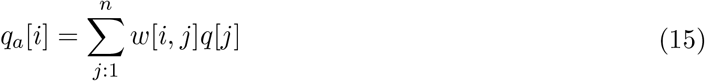

Expressing this in matrix form yields:

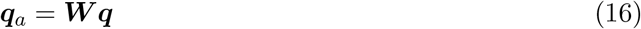

Note that, without loss of generality, we assume that the weights are scaled such that they add up to 1 for each kinase. This scaling ensures that the weights represent a weighted average. To estimate the standard error *σ*_*a*_[*i*] and the corresponding degrees of freedom *df*_*a*_[*i*] in the inferred activity *q*_*a*_[*i*] of kinase *i*, we use the Satterthwaite approximation:

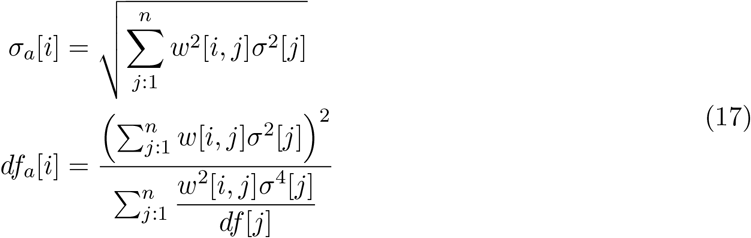

Expressing this in matrix form yields:

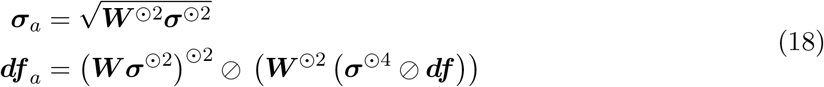

To clarify, the symbol ^⊙*k*^ represents the operation of taking the element-wise *k*th power (also known as the Hadamard power) of a matrix or vector. For example, ^⊙2^ corresponds to the element-wise square operation. Similarly, ⊘ represents element-wise division.

After ***q***_*a*_, ***σ***_*a*_ and ***df*** _*a*_ are estimated, to assess the statistical significance of a kinase *i*, a t-test is performed based on the t-statistic *t*_*a*_[*i*]:

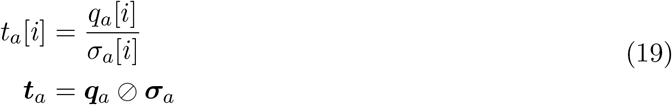

which follows a t-distribution with *df*_*a*_[*i*] degrees of freedom under the null hypothesis.

Note that, although the open formulas (such as in Equation (17)) are provided for clarity, in the implementation, their matrix correspondences (such as in Equation (18)) are performed using efficient sparse matrix operations for improved computational performance.

#### 2.6.3 Obtaining weights expressing the underlying formula for RoKAI inference

The RoKAI algorithm utilizes a heterogeneous network, denoted as 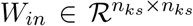, where the nodes represent kinases and/or phosphosites. The total number of nodes in the network is denoted as *n*_*ks*_ = *n*_*kin*_ + *n*, which is the sum of the number of kinases (*n*_*kin*_) and the number of phosphosites (*n*). The edges in this network capture various functional associations between kinases, phosphosites, and their combinations. To propagate the phosphorylation values across this functional network, the RoKAI algorithm employs an electric circuit model and solves a system of equations, as described in the reference [Yilmaz et al., 2021] and outlined below:

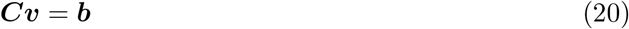

In the given equation, the matrix 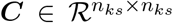 represents the conductance between nodes in the network, allowing a portion of phosphorylation to be transfered to nearby nodes in the form of current. The vector 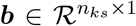 indicates the phosphorylation levels (log fold changes), while 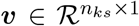 represents the node potentials to be computed. These node potentials ***v*** reflect the phosphorylation levels after the information has propagated through the network.

It is important to emphasize that not all nodes in the network are required to be quantified. Even nodes that do not have a computed fold change value, such as those with missing values in the experimental data, can still be retained in the network. In the case of an unquantified node denoted as *i*, its corresponding entry in the vector ***b*** is assigned a value of *b*[*i*] = 0 to indicate that it does not contribute to the fold change calculation and only kept as bridge node connecting other nodes.

The solution vector ***v*** of this system can be obtained efficiently using standard linear algebra solvers. However, in addition to finding the the solution vector ***v***, our goal here is to compute the weights 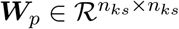 that represent the underlying solution of this system such that:

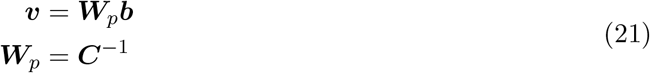

Obtaining the weighting matrix ***W***_*p*_ explicitly through the matrix inversion operation described above becomes computationally expensive, especially for typical large-scale networks with thousands or more nodes (phosphosites or kinases). This approach becomes even more challenging when working with limited computational resources, such as those available on a web server. To overcome this challenge, we have implemented several optimizations for the RoKAI algorithm to improve computational efficiency that involves computing a partial inverse. These optimizations ensure that the algorithm remains feasible and scalable, even for large-scale networks.

First, we introduce the concept of *relevant nodes*. These are phosphosites or kinases that have a functional annotation in the network, meaning they are associated with at least one edge in the functional network ***W***_*in*_. We denote the number of relevant nodes as *n*_*rel*_, and *I*_*rel*_ represents the indices of these nodes. To focus on the relevant nodes, we define 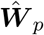 as the subset of the network ***W***_*p*_ that only includes the relevant nodes, represented by the indices *I*_*rel*_. In other words, 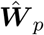 corresponds to the submatrix ***W***_***p***_[*I*_*rel*_, *I*_*rel*_], which has a size of *n*_*rel*_ *× n*_*rel*_. Similarly, we ***Ĉ*** define as the subset of the conductance matrix that corresponds to the relevant nodes.

In addition to the concept of relevant nodes, we introduce the notion of *quantified nodes*. These nodes refer to the phosphosites that are explicitly identified in the dataset being processed. In other words, they are phosphosites for which a fold change value is computed (i.e., does not have a missing value). In general, we are only interested in computing the further subset of 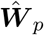 that involves the quantified nodes. We refer to this subset as 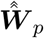 which is of size *n*_*q*_ *× n*_*q*_ where *n*_*q*_ is the number of nodes that are both quantified and relevant. The remaining nodes that are relevant but not quantified play a crucial role as bridges connecting other nodes in the network. While these are necessary for obtaining the solution of the system as they contribute to the overall connectivity and information flow, they need not be explicitly expressed in the final solution or underlying formula behind the inferred kinase activities.

We can partition the conductance matrix ***Ĉ*** into 2 *×* 2 blocks based on on the quantification status of the nodes:

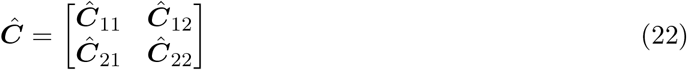

In this partitioning scheme, we assign a label of “1” to the block corresponding to the quantified nodes, indicating their quantification status, and a label of “2” to the block representing the remaining nodes. Similarly, we can express the linear system in Equation (20) using the partioning as follows:

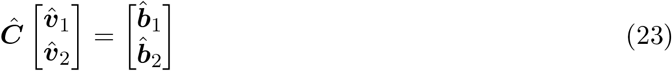

Since the unquantified nodes in block “2” does not have computed fold changes, 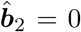 and the above equation can be rewritten as:

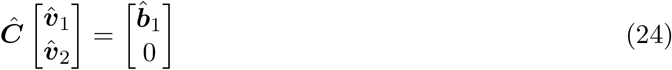

What we are looking for here is a matrix 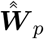, a partial inverse of ***Ĉ*** that will satisfy:

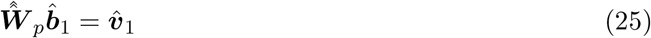

Given that the block matrix ***Ĉ***_11_ is invertible, we can obtain a partial inverse of matrix ***Ĉ*** inverting ***Ĉ***_11_ and replacing the corresponding block ***Ĉ***_22_ with the Schur complement ***Ĉ****/****Ĉ***_11_ and adjusting the off-diagonal elements of the resulting matrix accordingly [Tsatsomeros, 2000]:

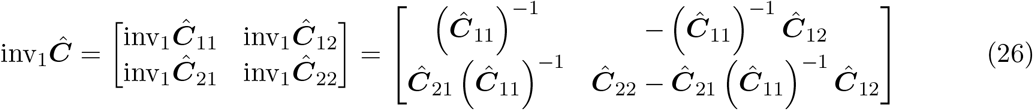

This partial inversion corresponds to a rotation of the matrix and satisfies the following property [Tsatsomeros, 2000]:

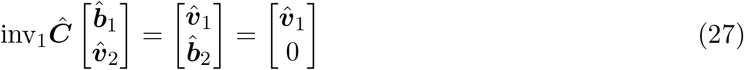

Thus, this produces two main equations:

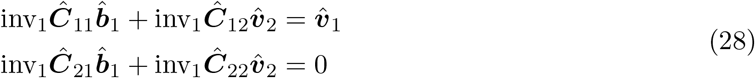

Reorganizing the second equation above yields:

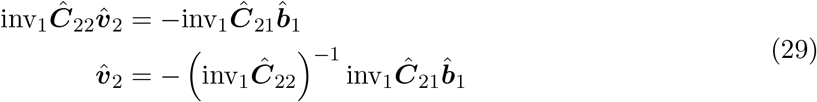

Substituting 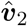 into the first equation results in:

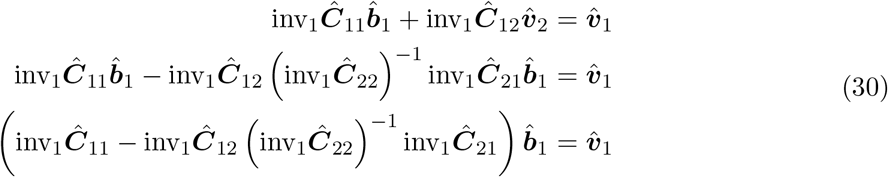

Thus,

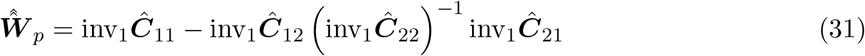

Substituting the values for the entries of the partial inverse, inv_1_***Ĉ*** matrix results in:

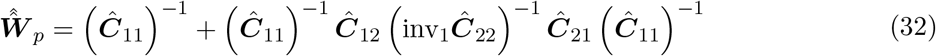

Here, the computation of the inverse (inv_1_***Ĉ***_22_)^−1^ is still computationally costly. Fortunately, we do not have to explicitly compute it. The multiplication ***Ĉ***_12_ (inv_1_***Ĉ*** _22_)^*−1*^ corresponds to the solution ***S*** of the following linear system, which can be efficiently solved (e.g., using *mrdivide* function in Matlab or *solve* function in R):

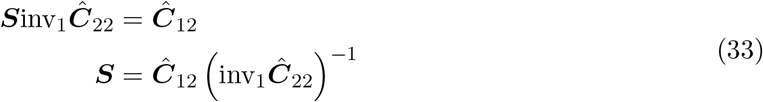

where

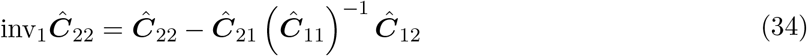

Substituting the solution matrix *S* yields the equation:

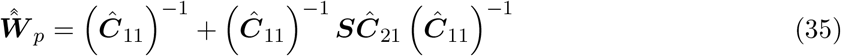

Note that, the weighting matrix 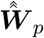 of size *n*_*q*_ *× n*_*q*_ is defined for quantified and relevant nodes. However, it can be easily mapped to the space of all phosphosites ***W***_*p*_ (of size *n × n*) by setting the remaining values for all other nodes as 0.

After the weighting matrix for phosphosites ***W***_*p*_ is obtained, the weights kinase inference ***W*** are obtained for mean substrate phosphorylation using the kinase-substrate network ***W***_*ks*_:

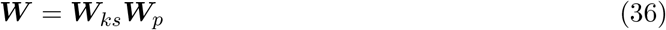

Here, the matrix ***W*** represents a weighted network of size *n*_*kin*_ *× n*, where *n*_*kin*_ denotes the number of kinases and *n* represents the number of phosphosites. On the other hand, the matrix ***W***_*ks*_ represents an unweighted network of size *n*_*kin*_ *× n*, specifically indicating the interactions between kinases and phosphosites that are known kinase targets.

#### 2.6.4 Options for kinase activity inference

RokaiXplorer offers several options specific for kinase activity inference. These are outlined below:

##### Kinase-substrate dataset

The options are PhosphoSitePlus (PSP) or PSP + Signor. The former only uses the known kinase targets *W*_*ks*_ from PSP [Hornbeck et al., 2015], and the latter additionally include known targets from Signor [Licata et al., 2020] as well.

##### RoKAI Network

Determines what kind of interactions are included in the RoKAI functional network *W*_*in*_. *KinaseSubstrate* option only includes the known kinase targets *W*_*ks*_ network. *KS+PPI* option additionally adds the protein-protein interactions (PPI) between the kinases. *KS+PPI+SD* also includes interactions between phosphosites based on structure distance evidence. *KS+PPI+SD+CoEv* further includes interactions between phosphosites based on co-evolution evidence.

##### Use sites in functional network

A binary flag that determines whether the network propagation through RoKAI should be performed or not.

##### Min. number of substrates

Determines the minimum number of phosphosites that are known targets of a kinase needs to be identified in the dataset for that kinase to be included in the analysis. In other words, the kinase will be considered only if it has at least the specified minimum number of identified phosphosites in *W*_*ks*_.

### 2.7 Statistical inference at pathway level

In addition, RokaiXplorer provides the functionality to perform pathway or gene ontology (GO) term enrichment analysis based on peptides or proteins that have been identified as significant in previous analyses involving phosphorylation or protein expression. Pathway analysis is a valuable tool for gaining insights into the biological pathways and networks affected by the observed changes in the phosphoproteome. By conducting pathway analysis, users can further understand the broader biological context and functional implications of the identified phosphorylation events.

The enrichment analysis in RokaiXplorer is conducted using an over-representation analysis (ORA) approach. This involves assessing the enrichment of significant phosphosites or proteins in specific pathways or gene ontology terms. It is important to note that the analysis is performed at the protein level, even when examining phosphorylation at the phosphosite level. Specifically, for enrichment analysis on phosphosites, we map the significant phosphosites to their corresponding proteins. A protein is considered significant if it contains at least one significant phosphosite. To determine the statistical significance of the enrichment, we employ the chi-squared test with Yate’s correction [Yates, 1934]. This test helps evaluate whether the observed distribution of significant phosphosites or proteins across pathways or gene ontology terms deviates significantly from what would be expected by chance alone and produces the p-values. The test produces p-values, which indicate the strength of evidence for enrichment. We apply the Benjamini-Hochberg procedure [Benjamini and Hochberg, 1995] to alleviate the multiple comparisons issue by limit the false discovery rate (FDR) of the findings.

#### 2.7.1 Estimating magnitude of enrichment — Bayesian log risk ratio

To estimate the magnitude of enrichment, RokaiXplorer utilizes a Bayesian approach to estimate log risk ratio. These estimates are primarily used to visualize the magnitude of enrichment, such as in bar plots in the inspection window. Let 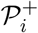 denote the set of significant proteins within pathway *i* (e.g., the *hits*), and 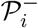 denote the set of non-significant proteins within the pathway (i.e., the *misses*). Similarly, 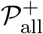 represents the set of all significant proteins, and 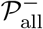 represents the set of all non-significant proteins. Furthermore, *n*^+^[*i*] corresponds to the number of significant proteins in pathway *i*, while *n*^*−*^[*i*] corresponds to the number of non-significant proteins within the pathway. Similarly, 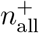 denotes the total number of significant proteins, and 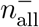 denotes the total number of non-significant proteins. Without using a Bayesian prior, the log risk ratio (logRR) can be computed as follows:

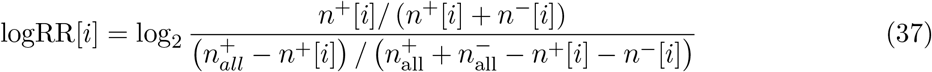

This estimate presents several issues. For instance, when a pathway contains only 1 significant protein (1 hit and 0 misses), the risk ratio approaches the maximum value possible (equal to 1 divided by the significance ratio 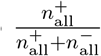), even though the presence of only 1 significant protein provides weak evidence that cannot be reliably attributed to anything other than chance alone. In contrast, a pathway with 100 significant proteins out of 100 proteins yields a similar risk ratio, despite providing much stronger and more confident evidence. To address this issue, we employ a Bayesian estimate that incorporates a prior belief, assuming that the pathway’s enrichment (hit ratio) is equal to the significance ratio 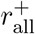 when no additional evidence is available:

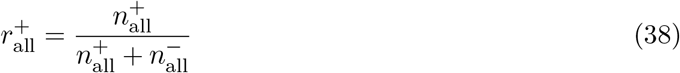

For this purpose, our perspective is similar to that of the coin flip problem. Imagine that we conduct a series of independent trials, where each trial can result in a hit or a miss. After performing these trials, we observe a total of *n*^+^ hits and *n*^*−*^ misses. Now, our goal is to determine the likelihood that the coin is fair, or in other words, what is our posterior belief about the probability of getting a hit, denoted as *r*?

Assuming uniform prior for all possible *r* values, the answer lies in the beta distribution:

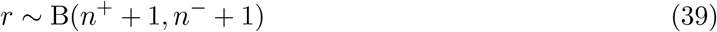

Here, we can generalize this approach by introducing *α*^+^ and *α*^*−*^, which represent the “prior number of hits” and “prior number of misses” respectively. These values act as pseudotrials and represents the prior belief in the estimation process:

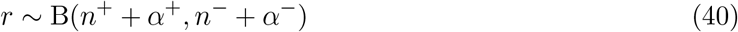

where 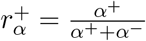 represents the prior mean. To select appropriate prior values, we consider three principles. First, we aim to ensure that 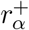 matches the population significance ratio 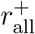. Second, we want *α*^+^ and *α*^*−*^ to be at least equal to 1 or higher. Lastly, we want the prior to be equivalent to uniform prior when 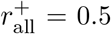 (where hits and misses are equally likely). To incorporate these principles, we utilize the following prior values:

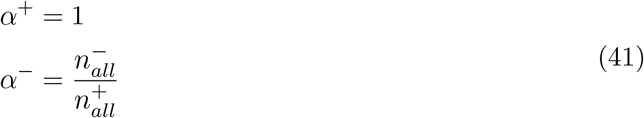

It is important to note that these prior values are chosen under the assumption that the number of significant proteins 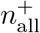 is smaller than the number of non-significant proteins 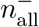. However, in the unlikely scenario where 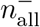 is larger than 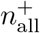, we swap the two groups and set 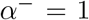 and 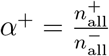 to satisfy the desired criteria and ensure appropriate estimation.

Based on these parameters, we calculate the medians *m*[*i*] and *m*_out_[*i*] of the posterior distribution for the hit ratios *r*[*i*] and *r*_out_[*i*] respectively. The hit ratio *r*[*i*] represents the ratio of significant proteins to all proteins in pathway *i*, while the hit ratio *r*_out_[*i*] represents the ratio of significant proteins to all proteins outside of pathway *i*. The medians *m*[*i*] and *m*_out_[*i*] are obtained based on the quantile function for beta distribution:

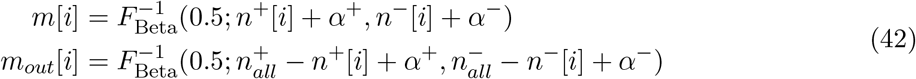

where 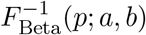 denotes the quantile function of the beta distribution for quantile *p* and parameters *a* and *b*. Thus, based on the provided parameters and Bayesian prior, we derive an estimate of the log risk ratio 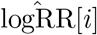 for pathway *i* as follows:

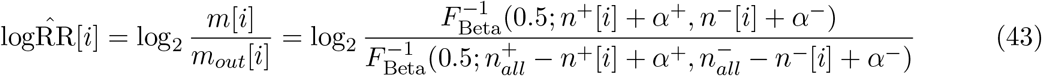

#### 2.7.2 Parameters for pathway enrichment

RokaiXplorer offers several options for the pathway enrichment. These are outlined below:

##### Inclusion criteria (enrichment terms)

These options determine the inclusion of enrichment terms in the analysis. The first option categorizes the terms based on their category, such as *Biological Process, Cellular Process*, or *Molecular Function* for GO terms. Additionally, there are options to filter out pathways based on the number of proteins identified in the input dataset. This can be done either by specifying a minimum number of observed proteins or by setting a minimum ratio of observed proteins to the total number of proteins in the pathway. Furthermore, there is an option to filter out highly similar pathways based on the Jaccard index. Pathways that exhibit a highly similar set of observed proteins are filtered out, retaining only one of them. In cases of duplication, the smaller pathway is retained Furthermore, there is an option to filter out highly similar pathways based on the Jaccard index. When this option is enabled, if multiple pathways exhibit a highly similar set of observed proteins, only one of them is retained, prioritizing the smaller pathway with less number of proteins.

##### Background set (proteins)

These options determine the set of significant proteins to be used for the enrichment analysis. The first option allows for selecting the data source, which can be either the results from the *Phosphosite, Phosphoprotein*, or *Protein expression* analyses. If the *Phosphosites* option is chosen, the background set will include proteins that have at least one significant phosphosite. The subsequent options define the criteria for determining significance, such as the cut-off based on p-values or log2 fold changes. Additionally, there is a binary setting to indicate whether the Benjamini-Hochberg procedure should be applied to control the false discovery rate (FDR) of the findings. Finally, an additional option is provided to restrict the set of significant proteins to either positive or negative log fold changes, if desired.

### 2.8 Inspection window: Performing sample-wise inferences

In addition to the specific statistical analyses described earlier (e.g., phosphosites, phosphoproteins, protein expression, kinase inference, and pathway enrichment), RokaiXplorer offers sample-wise analyses at the individual sample level for more detailed inspection in the inspection window. These sample-wise analyses compare each sample in the Case group to all samples in the Control group, providing granular results that capture the variance between samples. The inspection window allows for visualization of these results through bar plots or box plots, providing a comprehensive view of the individual sample-level analysis.

For the sample-wise analysis at the phosphosite and protein expression level, we utilize the z-test as a simpler alternative to the t-test. The z-test estimates the standard error by considering the variance across the phosphosites or proteins. Unlike the t-test, which requires multiple samples to measure the variance between them, the z-test can be applied without such a requirement. Furthermore, in the kinase activity inference, we perform the analysis without utilizing network propagation through RoKAI’s functional network. This decision is made to conserve the computa+tional resources on the web server and streamline the analysis process.

## Footnotes

1 https://github.com/serhan-yilmaz/RokaiXplorer

## References

Benjamini, Y. and Hochberg, Y. (1995). Controlling the false discovery rate: a practical and powerful approach to multiple testing. Journal of the Royal statistical society: series B (Methodological), 57(1):289–300.

Hornbeck, P. V., Zhang, B., Murray, B., Kornhauser, J. M., Latham, V., and Skrzypek, E. (2015). Phosphositeplus, 2014: pmutations, ptms and recalibrations. Nucleic acids research, 43(D1):D512–D520.

Licata, L., Lo Surdo, P., Iannuccelli, M., Palma, A., Micarelli, E., Perfetto, L., Peluso, D., Calderone, A., Castagnoli, L., and Cesareni, G. (2020). Signor 2.0, the signaling network open resource 2.0: 2019 update. Nucleic acids research, 48(D1):D504–D510.

Ritchie, M. E., Phipson, B., Wu, D., Hu, Y., Law, C. W., Shi, W., and Smyth, G. K. (2015). limma powers differential expression analyses for rna-sequencing and microarray studies. Nucleic acids research, 43(7):e47–e47.

Satterthwaite, F. E. (1946). An approximate distribution of estimates of variance components. Biometrics bulletin, 2(6):110–114.

Smyth, G. K. (2004). Linear models and empirical bayes methods for assessing differential expression in microarray experiments. Statistical applications in genetics and molecular biology, 3(1).

Tsatsomeros, M. J. (2000). Principal pivot transforms: properties and applications. Linear Algebra and its Applications, 307(1-3):151–165.

Yates, F. (1934). Contingency tables involving small numbers and the χ 2 test. Supplement to the Journal of the Royal Statistical Society, 1(2):217–235.

Yilmaz, S., Ayati, M., Schlatzer, D., C icek, A. E., Chance, M. R., and Koyutürk, M. (2021). Robust inference of kinase activity using functional networks. Nature communications, 12(1):1177.

Yilmaz, S., Lopes, F. B. T. P., Schlatzer, D., Wang, R., Qi, X., Koyuturk, M., and Chance, M. (2023). Exploring temporal and sex-linked dysregulation in alzheimer’s disease phosphoproteome. bioRxiv.

